# Activation of Endogenous Spinal Cord Stem Cell Mediates Recovery Induced by Olfactory Ensheathing Cell Transplantation Following Spinal Cord Injury

**DOI:** 10.1101/2025.07.22.666088

**Authors:** Quentin Delarue, Axel Honore, Tingting Xu, Chaima Chalfouh, Marine Di Giovanni, David Vaudry, Xiaofei Li, Nicolas Guérout

**Author notes:** Address correspondence to: Corresponding authors., Lead contact: Nicolas Guérout; Co-correspondance: Xiaofei Li, Co-correspondance: Quentin Delarue. Equal contribution (Xiaofei Li, Nicolas Guérout).

## Abstract

Spinal cord injury (SCI) leads to irreversible motor and sensory deficits and currently has no curative treatment. Among the various therapeutic strategies explored, cell transplantation— using either stem or differentiated cells—has been extensively studied. One of the most promising approaches involves the use of olfactory ensheathing cells (OECs), which have demonstrated unique potential to promote functional recovery and tissue repair following SCI. However, the mechanisms underlying these effects remain poorly understood.

In this study, we investigated how OEC transplantation modulates endogenous spinal stem cells, with a particular focus on ependymal cells. Using inducible transgenic mouse lines and fate-mapping approaches, we show that OEC transplantation significantly enhances ependymal cell proliferation and self-renewal both *in vivo* and *in vitro*. Moreover, OECs promote astrocytic differentiation of ependymal progeny, which express low levels of inhibitory molecules— suggesting a supportive role in creating a permissive scar microenvironment.

Transcriptomic analyses further revealed that OEC transplantation downregulates genes associated with axonal growth inhibition, thereby contributing to improved neuronal survival. Finally, by using a unique transgenic mouse model in which ependymal cell proliferation is genetically blocked, we demonstrate that the beneficial effects of OEC transplantation depend on ependymal cell activation.

Together, these findings establish ependymal cells as essential mediators of the regenerative response induced by OECs. They also highlight a therapeutic strategy based on activating and modulating endogenous stem cells via the transient presence of non-integrating transplanted glial cells. This work contributes to our understanding of SCI repair and supports the clinical potential of OEC-based cell therapies.

## INTRODUCTION

Spinal cord injury (SCI) remains a chronic, currently incurable condition, with incidence rates ranging from 10 to 246 cases per million people annually, depending on the geographic region studied (Golestani et al., 2022; Siddiqui et al., 2015). SCI commonly leads to irreversible deficits in motor and sensory functions below the site of injury and is frequently associated with complications such as neuropathic pain, spasticity, and autonomic dysfunctions (Soriano et al., 2023). At present, no treatment exists to promote functional recovery. Current medical management is limited to surgical intervention aimed at preventing secondary damage, and to symptomatic care addressing pain and infections during both the acute and chronic phases (MacLean and Okonkwo, 2025).

Given the complexity and progressive nature of cellular and functional deterioration following SCI, effective therapeutic strategies will likely require a combinatorial approach. Several interrelated mechanisms contribute to the failure of spontaneous regeneration and functional recovery, including the formation of a fibroglial inhibitory environment, neuronal loss, demyelination, axonal degeneration, limited axonal regrowth, and chronic inflammation (Clifford et al., 2023; Dias et al., 2021; Guérout, 2021). In this context, cell transplantation has emerged as a promising strategy to counteract cellular loss and reshape the inhibitory microenvironment (Sykova et al., 2021). Various cell types have been explored as potential biotherapies in preclinical models of SCI. Among them, olfactory ensheathing cells (OECs) have demonstrated the ability to promote functional recovery, making them a particularly attractive candidate for therapeutic development (Chen et al., 2024; Phelps et al., 2025; Ursavas et al., 2021). While the precise mechanisms remain unclear, it has been demonstrated that OECs secrete neurotrophic factors, modulate the extracellular matrix, but also help regulate the inflammatory response even on distant organs and contribute to the phagocytosis of debris in the injured area (Delarue et al., 2025; Delarue and Guérout, 2022; Murtaza et al., 2022). These unique properties have naturally led researchers to investigate the effects of OEC transplantation in the context of SCI. One of the first studies, although widely debated, reported that OEC transplantation applied to a paraplegic patient in a clinical trial produced remarkably beneficial effects (Tabakow et al., 2014, 2013). However, due to the lack of precise mechanisms of the contribution of endogenous spinal cord cells in response to the therapy, even though other patients that received similar OECs transplantation also achieved significant recovery, their recoveries did not reach the same level of the patient previously reported (Gómez et al., 2018). It is important to highlight that one of the advantages of OEC transplantation lies in the fact that these cells are already differentiated glial cells and do not persist long-term after transplantation (Khankan et al., 2016). As a result, they are incapable of differentiating into other cell types and pose no risk of teratoma formation. Therefore, to obtain better therapeutic potential from OECs for SCI, it is essential to further study how OEC transplantation could promote regeneration.

While the adult spinal cord has limited regenerative potential after SCI, previous fate-mapping studies using transgenic mouse models based on CreER driven by FoxJ1 promoter reveal that ependymal cells are the spinal cord stem cells and can give rise to low reactive astrocytes and oligodendrocytes after SCI as early as the juvenile stage across life span (Barnabé-Heider et al., 2010; Li et al., 2018, 2016; Meletis et al., 2008; Ripoll et al., 2023; Rodrigo Albors et al., 2023; Stenudd et al., 2022). Moreover, it has been shown that using a unique transgenic mouse model in which ependymal cell proliferation is specifically blocked upon SCI, ependymal cells play a critical role in glial scar formation and are required for neuronal survival adjacent to the lesion (Sabelström et al., 2013). This study confirms that ependymal cells are required for glial scar formation and to maintain the integrity of the injured spinal cord (Sabelström et al., 2013).

On the other hand, it has been shown that OEC transplantation can enhance axonal regrowth by reducing the severity of the glial scar, in which astrocytes exhibit lower levels of GFAP expression (Mayeur et al., 2013). These two observations suggest that OECs could modulate endogenous stem cells and their progeny after SCI, leading to a less reactive glial scar and higher regenerative potential of the injured spinal cord. To explore this hypothesis, we took advantage of transgenic mouse lines to specifically fate map ependymal cells in the presence of transplanted OECs both *in vivo* and *in vitro* after SCI, as well as characterize the regeneration after OEC transplantation with or without proliferative ependymal cells *in vivo*. Our study has demonstrated that OECs can promote neuronal survival and glial cell regeneration, but all these effects are highly dependent on ependymal cells, which serves as the central mediator.

## MATERIALS AND METHODS

### Animal experimentation

#### Animal care and use statement

The experimental protocol was designed to minimize pain and discomfort for the animals. All experimental procedures adhered to the European Community guidelines on the care and use of animals (86/609/CEE; Official Journal of the European Communities no. L358; 18 December 1986), the French Decree no. 97/748 of 19 October 1987 (Journal Officiel de la République Française; 20 October 1987), and the recommendations of the Cenomexa Ethics Committee.

Mice were housed in conventional, secure rodent facilities (two to five animals per cage, sexes separated) under a 12-h light/dark cycle with ad libitum access to food and water. In addition to WT C57BL/6, we used three transgenic mouse lines in this study:

- Adult mice tamoxifen-inducible FoxJ1-CreER^T2^(Jacquet et al., 2011; Jacquet et al., 2009; Ostrowski et al., 2003), with a C57BL/6 genetic background crossed with Rosa26-YFP (Figures 1-3) or Rosa26-TdTomato (Figure 4) reporter mice were used to fate map ependymal cells and their progeny.
- FoxJ1-CreER^T2^-Rasless-YFP (Rasless thereafter) (Figure 5) mice were used to conditionally delete the N-, K-, H-ras genes to block the proliferation of ependymal cells upon tamoxifen administration (Sabelstrom et al., 2013).
- Luciferase (LUX) line—CAG-luc-GFP (L2G85Chco+/+; FVB-Tg(CAG-luc,-GFP) L2G85Chco/J, JAX #008450) backcrossed to C57BL/6 for three generations to minimize strain-related variability were used to assess bOEC survival after transplantation.

**Figure 1.**
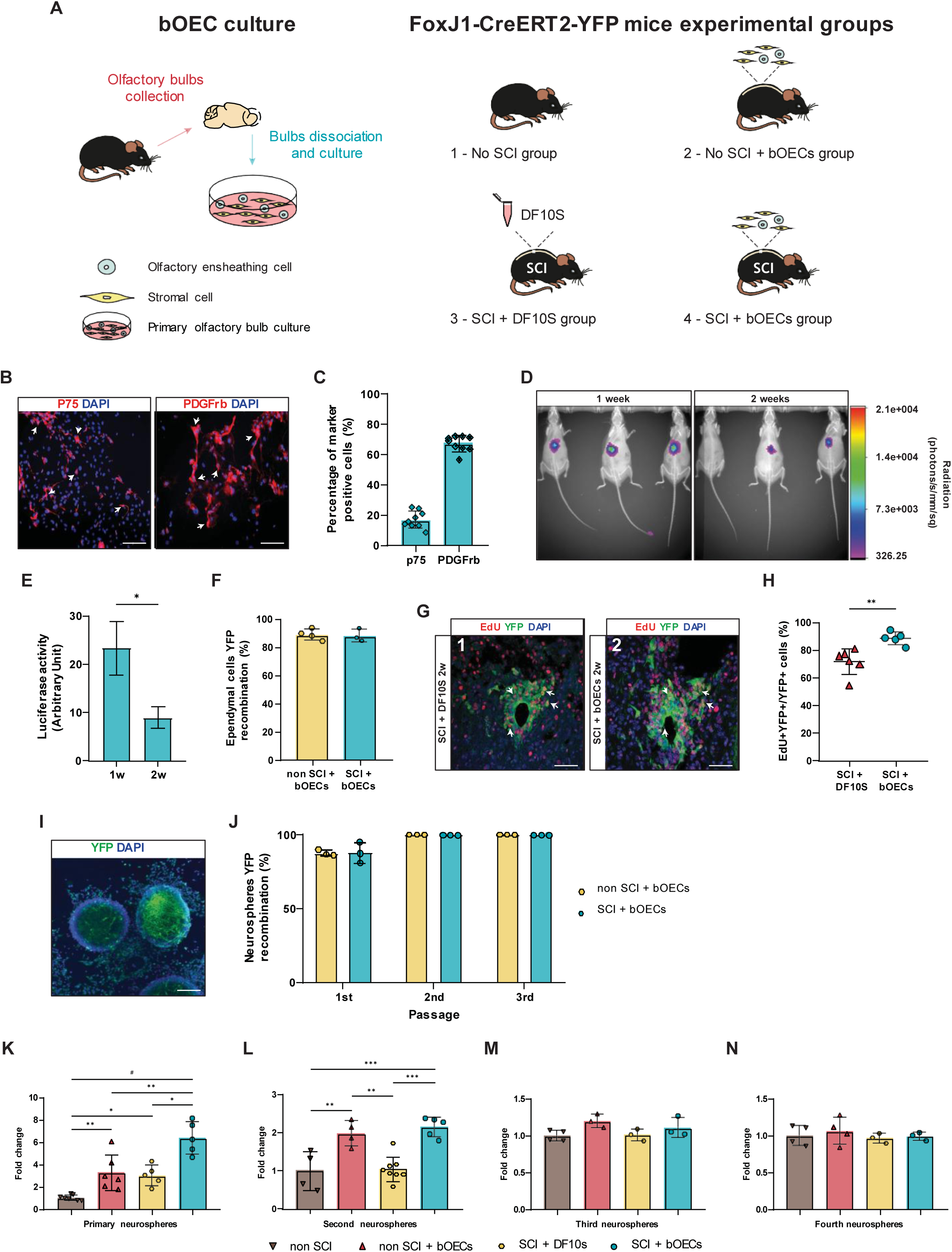
The transplantation of bOECs increases self-renewal capacity and proliferation of ependymal cells. **A**) Schematic representation of the four main experimental groups used in this section of our study. (1) No SCI group, uninjured animals used to define the baseline of *in vivo* and *in vitro* experiments. (2) No SCI + bOECs group, uninjured animals with bOEC transplantation. (3) SCI + DF10S group, these animals underwent SCI and DF10S culture medium was injected immediately. (4) SCI + bOECs group, these animals underwent SCI and primary bOECs were transplanted immediately. **B**) Representative pictures of bOEC culture characterization. Cells were stained with anti-p75 and anti-PDGFRβ antibodies. Scale bar: 200 µm. **C**) Quantification of percentage of p75+ OECs and PDGFRβ+ fibroblasts in bOEC cultures. **D**) Representative images of luciferase+ cells in WT mice 1 week and 2 weeks post-transplantation. **E**) Quantification of luciferase signals in WT mice 1 week and 2 weeks post-SCI and transplantation. **F**) Quantification of *in vivo* YFP recombined ependymal cells from FoxJ1-CreER^T2^-YFP mice after bOEC transplantation from non-SCI and SCI mice. **G**) Representative images of coronal section of adult lesioned spinal cord from FoxJ1-CreER^T2^-YFP mice 2 weeks after SCI with DF10S injection (**G1**) and SCI with bOEC transplantation (**G2**). Sections were stained with Edu Click-it©. Scale bar: 25µm. **H**) Quantification of proliferating EdU+YFP+ positive recombined cells *in vivo* 2 weeks after SCI. **I**) Representative image of primary cultured recombined neurospheres from FoxJ1-CreER^T2^-YFP injured mice. Scale bar: 200 µm. **J**) Comparative quantification of recombined neurospheres over passages from FoxJ1-CreER^T2^-YFP mice after bOEC transplantation from non-SCI and SCI mice. **K-N**) Quantification of the total number of neurospheres derived from 100,000 cells over passages from SCI or uninjured WT mice that received DF10S or bOECs 1 week after injury. N=9 (**C**), 3 (**E**), 3-4 (**F**), 5-6 (**H**), 3 (**J**) and 3-7 (**K-N**) animals. Quantifications are expressed as average ± SD. Statistical evaluations were based on T-Test (**C**, **E**, **F**, **H**, **J** and **K**) and Kruskal-Wallis tests (**L-O**). * = P< 0.05, ** = P< 0.01 and *** = P< 0.001

**Figure 2.**
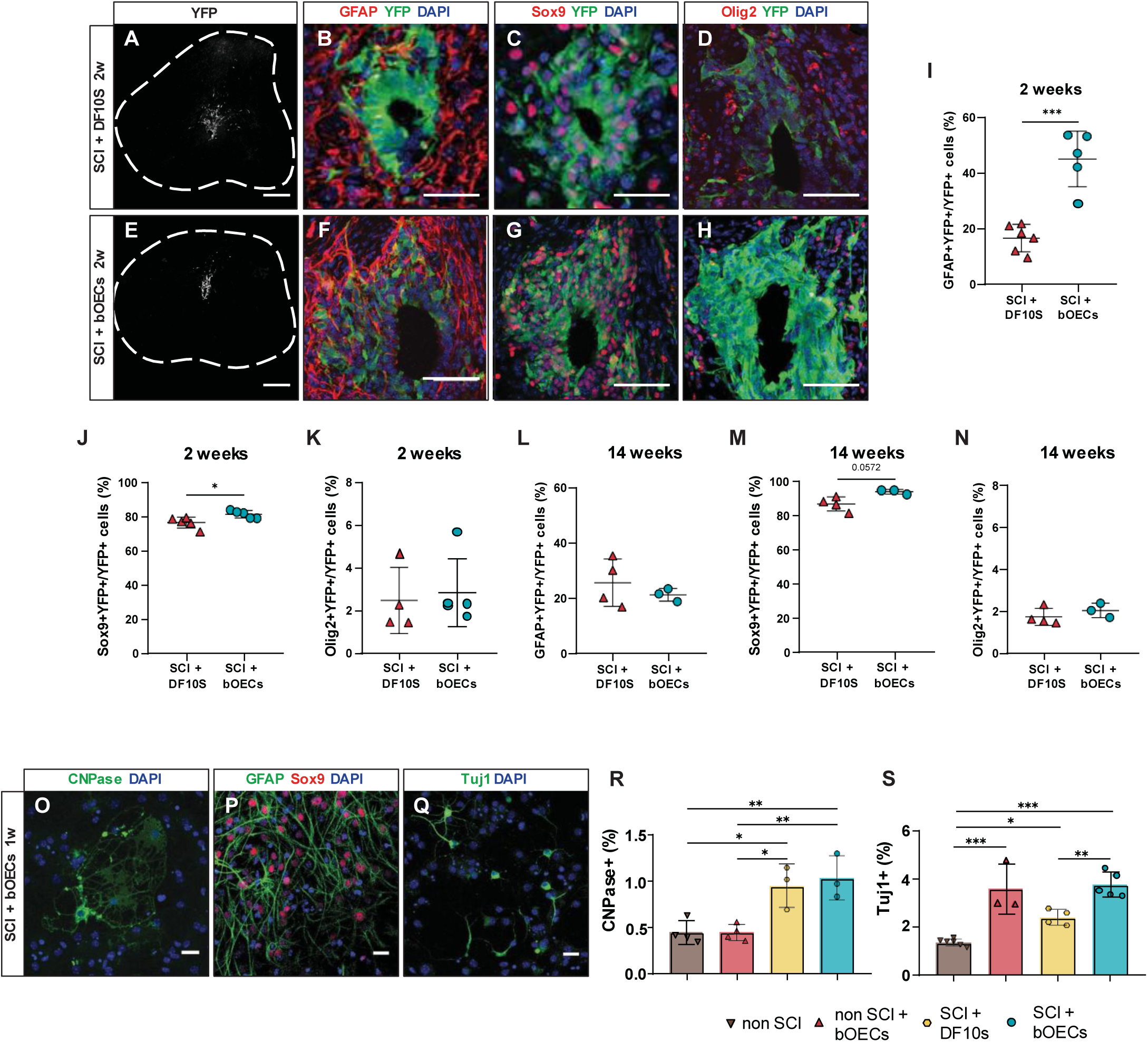
bOEC transplantation changes ependymal cell progeny *in vivo* and *in vitro*. **A-H**) Representative images of coronal sections of adult lesioned spinal cord from FoxJ1-CreER^T2^-YFP mice 2 weeks after SCI with injection of DF10S (**A-D**) and with transplantation of bOECs (**E-H**). Sections were stained with anti-GFAP, anti-Sox9 and anti-Olig2 antibodies. Scale bar: 200µm (**A** and **E**) and 50µm (**B**, **C**, **D**, **F**, **G** and **H**). Identification of cells derived from YFP+ recombined ependymal cells were identified by GFAP (**B** and **F**), Sox9 (**C** and **G**) and Olig2 (**D** and **H**). **I-N**) Quantification of double positive recombined cells in the spinal cord 2 weeks (**I-K**) and 14 weeks (**L-N**) after SCI with DF10S injection or bOEC transplantation. Quantification of YFP+GFAP+ cells (**I** and **L**). Quantification of YFP+Sox9+ cells (**J** and **M**). Quantification of YFP+Olig2+ cells (**K** and **N**). **O-Q**) Representative images of differentiated cells derived from primary neurospheres from WT adult mice one week after SCI and bOEC transplantation. Sections were stained with anti-CNPase (**O**), anti-Sox9 (**P**) and anti-Tuj1 (**Q**) antibodies. Scale bar: 25µm. **R**) Quantification of CNPase+ differentiated cells generated from primary neurospheres of adult WT mice from non-SCI, non-SCI + bOECs, SCI + DF10S and SCI + bOECs mice. **S**) Quantification of Tuj1+ differentiated cells generated from primary neurospheres of adult WT mice from non-SCI, non-SCI + bOECs, SCI + DF10S and SCI + bOECs mice. N=4-6 (**I-K**), 3-4 (**L-N**) and 3-6 (**R** and **S)** animals. Quantifications are expressed as average ± SD. Statistical evaluations were based on T-Test (**I-N**) and Kruskal-Wallis tests (**R** and **S**). * = P< 0.05, ** = P< 0.01 and *** = P< 0.001

**Figure 3.**
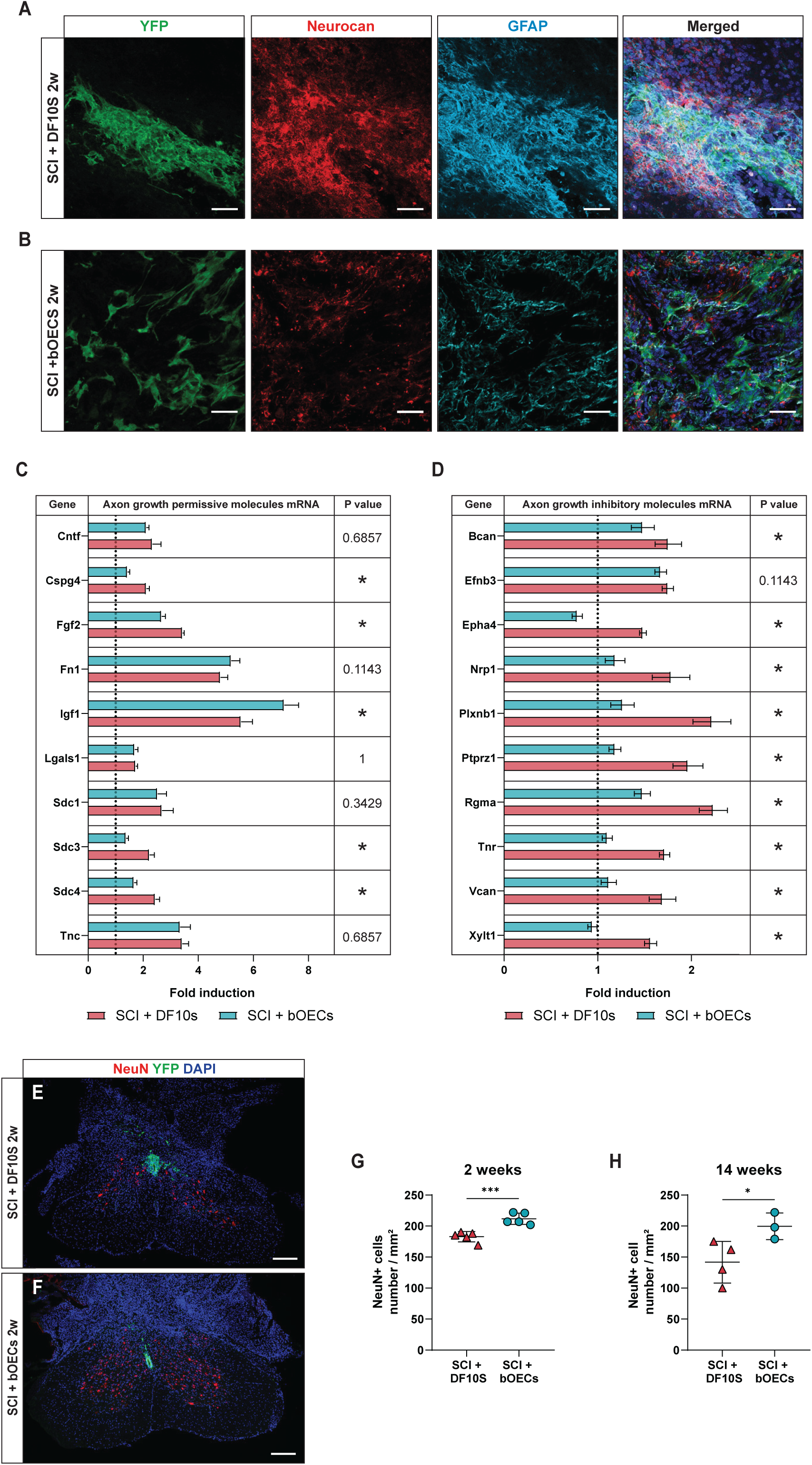
bOEC transplantation enriches the scar environment for axonal regrowth and promotes neuronal survival after SCI. **A-B**) Representative images of longitudinal section of adult injured spinal cord from FoxJ1-CreER^T2^-YFP mice 2 weeks after SCI with DF10S injection (**A**) and bOEC transplantation (**B**). Sections were stained with anti-GFAP and anti-Neurocan antibodies. Scale bar: 50µm. **C-D**) Histograms of mRNA expression of WT spinal cord of axon growth permissive molecules (**C**) and axon growth inhibitory molecules (**D**) 2 weeks after SCI with DF10S injection or bOEC transplantation. Dashed line corresponds to mRNA expression from uninjured control WT mice. **E-F**) Representative images of coronal sections of adult injured spinal cord from FoxJ1-CreER^T2^-YFP mice under SCI + DF10S (**E**) and SCI + bOECs (**F**) condition. Sections were stained with anti-NeuN antibody. Scale bar: 200µm. **G**) Quantification of the NeuN+ cells from WT mice 2 weeks after SCI with DF10S injection or bOEC transplantation. **H**) Quantification of the NeuN+ cells from WT mice 14 weeks after SCI with DF10S injection or bOEC transplantation. N=4 (**C** and **D**) and 3-5 (**G** and **H**) animals. Quantifications are expressed as average ± SD. Statistical evaluations were based on T-Test. * = P< 0.05, ** = P< 0.01 and *** = P< 0.001

**Figure 4.**
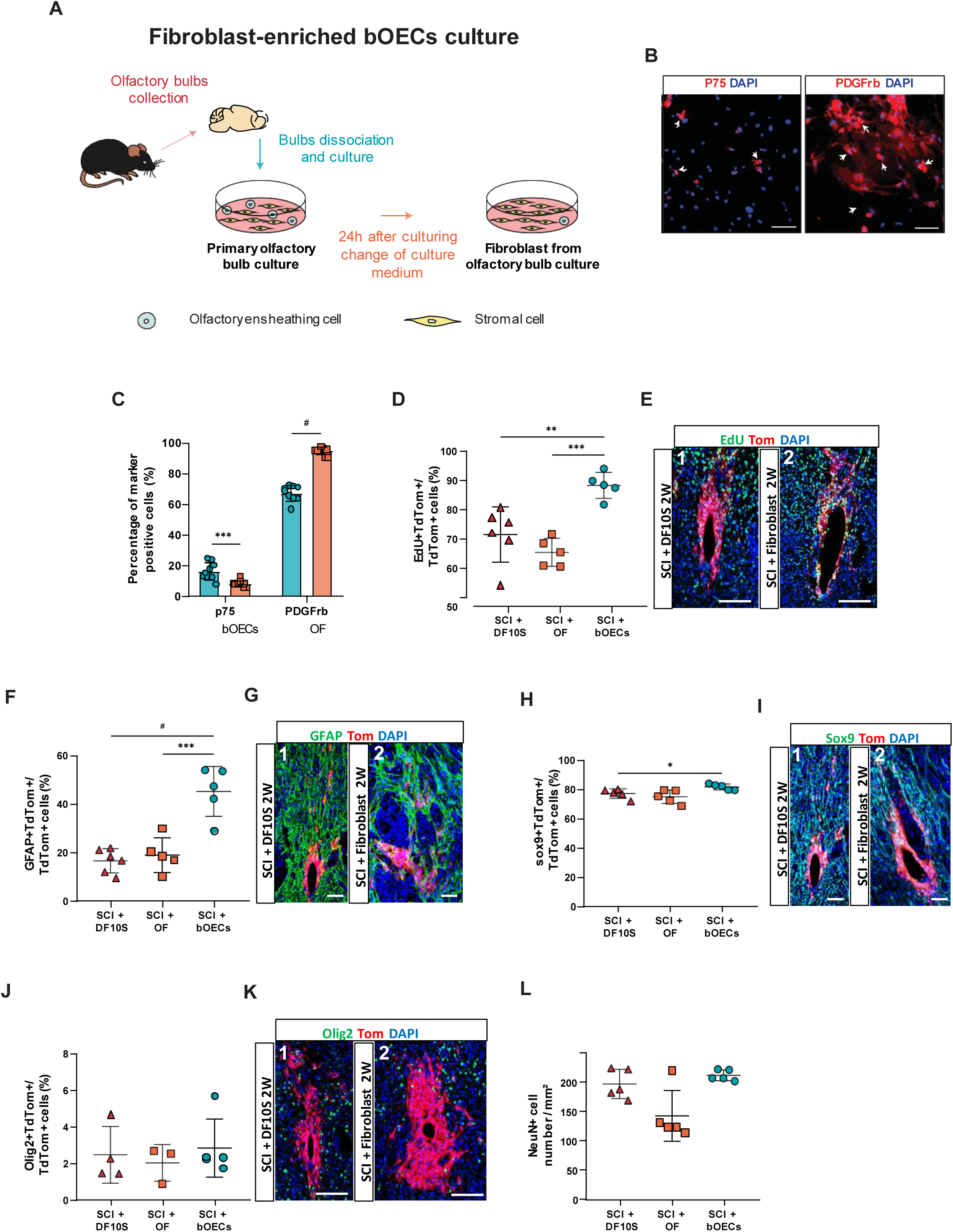
The effects of cell transplantation on ependymal cells are bOEC-dependent. **A**) Schematic representation of the culture method used to isolate olfactory fibroblasts. Twenty-four hours after plating, the supernatant is replaced to remove any bOECs that have not yet adhered to the plastic dish. **B**) Representative pictures of olfactory fibroblast culture characterization. Sections were stained with anti-p75 and anti-PDGFRβ antibodies. Scale bar: 200 µm. **C**) Quantification of percentage of p75+ OECs and PDGFRβ+ fibroblasts. **D**) Quantification of proliferating EdU+TdTomato+ (TdTom) positive recombined cells *in vivo* 2 weeks after SCI. **E**) Representative images of coronal sections of adult lesioned spinal cord from FoxJ1-CreER^T2^-TdTomato mice 2 weeks after SCI with DF10S injection (**E1**) and SCI with bOEC transplantation (**E2**). Sections were stained with Edu Click-it©. Scale bar: 50µm. **F-K**) Quantification of double positive recombined cells in the spinal cord from FoxJ1-CreER^T2^-TdTomato mice 2 weeks after SCI with DF10S injection, fibroblast transplantation or bOEC transplantation. **G, I** and **K**) Representative images of coronal sections of adult lesioned spinal cord from FoxJ1-CreER^T2^-TdTomato mice 2 weeks after SCI with injection of DF10S (**G1, I1** and **K1**) and with transplantation of fibroblats (**G2, I2** and **K2**). Sections were stained with anti-GFAP (**G**), anti-Sox9 (**I**) and anti-Olig2 (**K**) antibodies. Scale bar: 50µm. **F**). Quantification of TdTomato+GFAP+ cells. **H**). Quantification of TdTomato+Sox9+ cells. **J**). Quantification of TdTomato+Olig2+ cells (**G**). **L**) Quantification of the NeuN+ cells from WT mice 2 weeks after SCI with DF10S injection, fibroblast transplantation or bOEC transplantation. N=8-9 (**C**) and 3-6 (**D, F, H, J** and **L**) animals. Quantifications are expressed as average ± SD. Statistical evaluations were based on T-Test (**C**) and Kruskal-Wallis tests (**D, F, H, J** and **L**). * = P< 0.05, ** = P< 0.01, *** = P< 0.001 and # = P< 0.0001.

**Figure 5.**
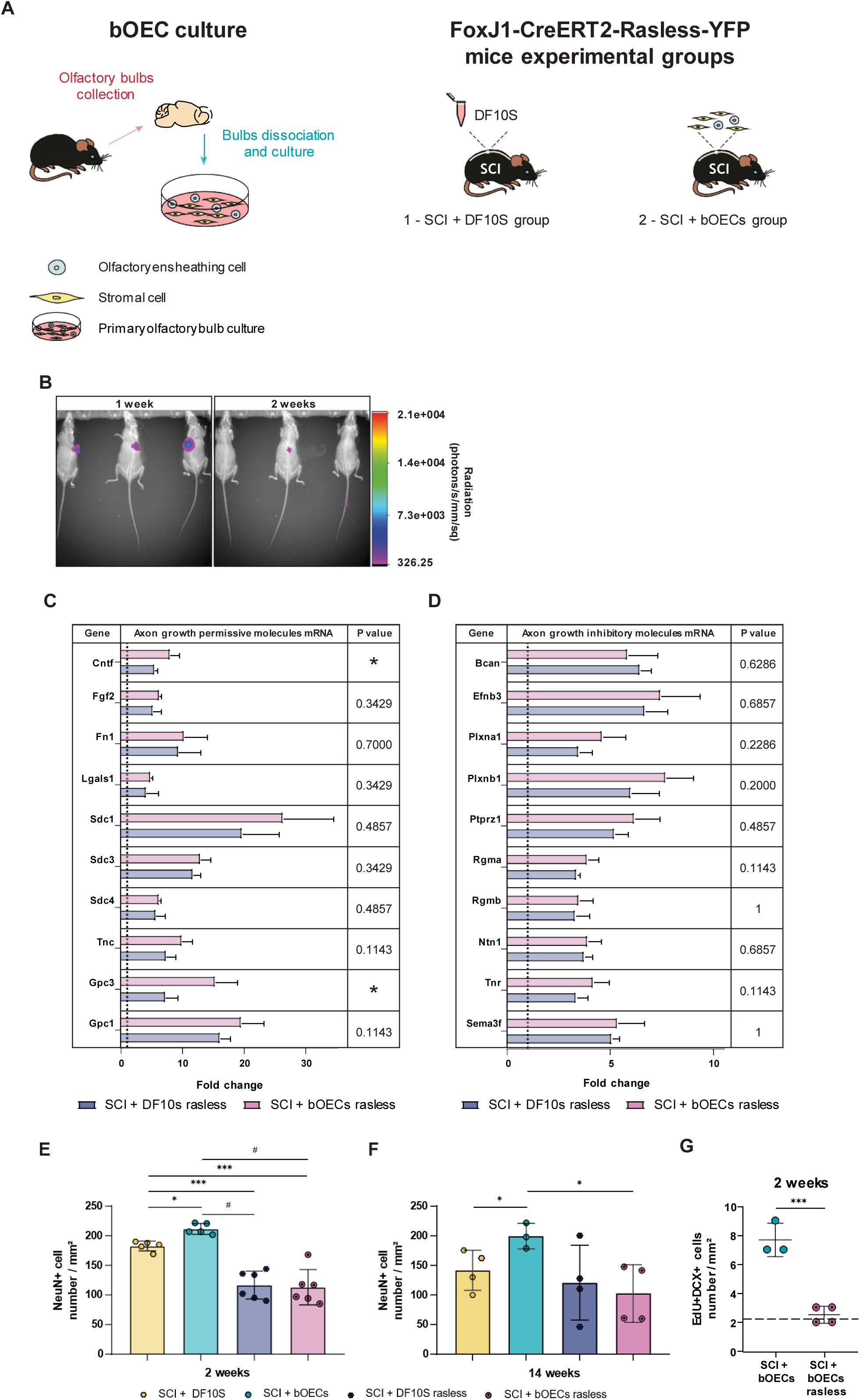
The endogenous ependymal cells are required for regenerative effects promoted by bOEC transplantation. **A**) Schematic representation of the two experimental groups used in this section of our study. In this section FoxJ1-CreER^T2^-YFP-Rasless mice were used to inhibit cell division in ependymal cells. (1) SCI + DF10S group, these animals underwent SCI and DF10S culture medium was injected immediately. (2) SCI + bOECs group, these animals underwent SCI and primary bOECs were transplanted immediately. **B**) Representative images of luciferase+ cells in FoxJ1-CreER^T2^-YFP-Rasless mice 1 week and 2 weeks post SCI and bOEC transplantation. **C-D**) Histograms of expression of FoxJ1-CreER^T2^-Rasless-YFP spinal cord of axonal growth permissive molecules (**C**) and axon-growth inhibitory molecules (**D**) after SCI with DF10S injection or bOEC transplantation. Dashed line corresponds to mRNA expression from uninjured control WT mice. **E-F**) Surviving NeuN+ cells in adult injured spinal cord from FoxJ1-CreER^T2^-YFP and FoxJ1-Rasless mice 2 weeks (**E**) and 14 weeks (**F**) post SCI with DF10S injection or bOEC transplantation. **G**) Quantification of the newborn neurons identified by EdU+DCX+ cells in adult injured spinal cord from FoxJ1-CreER^T2^-YFP and FoxJ1-Rasless mice 2 weeks post SCI with bOEC transplantation. Dash line represents the SCI + 10S group. N=4 (**C** and **D**), 3-6 (**E** and **F)** and 3-4 (**G**) animals. Quantifications are expressed as average ± SD. Statistical evaluations were based on T-test (**C, D and G**) and Kruskal-Wallis tests (**E** and **F).** * = P< 0.05, ** = P< 0.01, *** = P< 0.001 and # = P< 0.0001.

For both inducible mouse lines, to induce recombination, we injected 60 mg/kg of body weight once daily for 5 days. Clearance of tamoxifen was allowed for one week before the start of the SCI experiments.

Each experimental group was balanced for sex (50 % male / 50 % female) and main experimental groups are organized as follows (Figure 1A):

1. No SCI: unlesioned animals, providing baseline for *in vitro* and *in vivo* analyses.
2. No SCI + bOECs: unlesioned animals receiving bOEC transplantation.
3. SCI + DF10S: animals subjected to SCI without cell transplantation.
4. SCI + bOEC: SCI animals receiving bOEC transplantation.

#### Surgical procedure and cell transplantation

SCI was performed at the T9-T10 level as previously described (Delarue et al., 2019). After a midline dorsal incision and laminectomy of the T7 vertebra, a dorsal hemisection was performed with 25 Gauges needle. This lesion model induces the reactivity of endogenous stem cells without causing severe motor or functional impairments (Barnabé-Heider et al., 2010; Li et al., 2016; Sabelström et al., 2013). Immediately afterward, bOECs were transplanted with a micromanipulator-mounted stereotactic arm (World Precision Instruments). A sterile 1 mm glass capillary was lowered 1 mm deep, 1.5 mm lateral to the midline, 5 mm rostral and 5 mm caudal to the lesion. At each of the two sites, 2 µL of cell suspension (25 000 cells/µL) were injected over 1 min. The musculature and skin were sutured and animals were monitored daily; none of the mice developed skin lesions, infections, or autotomy during the study.

EdU, 5–ethynyl–2′–deoxyuridine (0,075mg/mL and 1% of sucrose, life) was administered with drinking water, exchanged every third day and kept in the dark. Following SCI, EdU was given twice by intraperitoneal injections (1,5mg/mL in PBS, 100µL per injection) at 6 hours interval, followed by administration in the drinking water for up to 7 days.

#### bOECs survival analysis

bOECs survival was assessed by bioluminescence. First, LUX+/− bOECs were prepared from OB of luciferase-reporter mice (see above for mouse line) and transplanted into wild-type or into Rasless recipients immediately after SCI. Bioluminescent emission from the grafted cells was captured with an *in-vivo* X-TREM 4XP cooled-CCD optical imager (Bruker). Mice received an intraperitoneal injection of D-luciferin (0.3 mg/g, XenoLight™, Perkin Elmer) and were imaged 30 minutes after injection at one and two weeks after SCI. Photon counts were quantified with Bruker molecular-imaging software as previously described (Takano et al., 2017).

### Primary olfactory bulb culture, purified olfactory fibroblasts cultures and cell transplant preparation

Olfactory bulb (OB) primary cultures were prepared as described previously by our team with slight modifications (Guerout et al., 2010; Honore et al., 2012). Mice were killed by lethal dose of sodium pentobarbital (150 mg/kg body weight) and decapitated. OB were immediately dissected and placed into Hank’s buffered salt solution (HBSS) after removing meninges. HBSS containing 0.1% of trypsin (Invitrogen) and OB were incubated for 20 min at 37°C. Trypsinization was stopped by adding Dulbecco’s Modified Eagle’s/Ham’s F12 medium (D.M.E.M/F12, Invitrogen), supplemented with 10% Fetal Bovine Serum (F.B.S, Invitrogen) and 0.5% penicillin/streptomycin (Invitrogen) (DF10S). The tissue was centrifuged at 1.500 RPM for 5 min, resuspended in DF10S and centrifuged again. Tissue was then triturated using a micropipette, until a homogenous cell suspension was obtained. Cells were plated in DF10S in 75 cm^2^ flasks (SARSTED). The flasks were incubated at 37°C, 5% CO_2_. The medium was changed every two days. Three to four weeks after plating, OB primary cultures (bOECs) were confluent.

Olfactory fibroblasts have been also generated from primary OB cultures. For that fibroblasts were selected using a purification method based on the one described by Nash et al. (Nash et al., 2001). To do so, OBs were cultured as described above, and on the following day, the plastic-adherent cells were collected and re-seeded into new culture dishes in DF10S for two to three weeks allowing to obtain purified cultures of fibroblasts (Figure 4A).

Before surgery, cultures were trypsinized to remove them from the dishes, and the cells were counted using a hemocytometer. bOEC or fibroblast culture cells were resuspended in DF10S at concentration of 25 000 cells/µL. DF10S was used as SCI control group (SCI + DF10S).

### Neural Stem Cells Cultures

Animals were sacrificed one week after SCI. Spinal cord cells were dissociated and neurosphere cultures were established as described (Li et al., 2016; Meletis et al., 2008). All cells isolated from one spinal cord were plated in 75 cm^2^ culture dishes. First, neurospheres were harvested after 2 weeks in culture and then were dissociated into single cells for passage or differentiation. Approximately 100,000 cells per animal were plated in a 25 cm^2^ culture dish for the next generation of neurospheres, and all the new neurospheres (second, third and fourth generations) were harvested after one week in culture. Dissociated primary neurospheres, approximately 70,000 cells/well, were plated in poly-L-lysine-coated chamber slides (Sigma) for differentiation with growth factors-free medium supplemented with 1% fetal bovine serum. Two to four independent experiments per group were performed. After 2 weeks in differentiation condition, immunocytochemistry was performed as described below.

### Immunocytochemistry

To characterize the primary bOEC or the fibroblast cultures (Figures 1A and 4A respectively), cells were detached from the culture dishes by trypsinization and subsequently seeded onto poly-L-lysine-coated chamber slides for two days. Then, the proportion of bOECs (P75-positive cells) and olfactory fibroblasts (PDGFrβ-positive cells) was quantified using immunocytochemistry experiments.

In order to assess the recombination rate after 14 days *in vitro* (Figure 1I and J), approximately 100–200 recombined primary neurospheres were plated in poly-L-lysine-coated chamber slides for one day, followed by fixation with 4% PFA (20 min at room temperature) and immunocytochemistry for quantification of recombined neurospheres was performed. For differentiation assay (Figure 2O-S), after 14 days in growth factor-free differentiation condition, immunocytochemistry was performed.

In all cases, one or two days after plating, the cells were washed with Phosphate Buffered Saline (PBS) and fixed in 4% paraformaldehyde (10 min at room temperature), the cells were washed with PBS and incubated in PBS/triton X100 0.1% (15 min at roomtemperature). Then, the cells were washed with PBS and incubated in PBS/BSA 2% containing primary antibodies overnight at 4 °C. After repeating washing with PBS, cells were incubated with the secondary antibodies conjugated antibodies. The wells were washed with PBS and incubated with Hoechst. Then, the cells were examined with a Leica microscope (Leica, Solms, Germany).

Full details of the primary antibodies used are reported in Table 1.

**Table 1.** Antibody table. Details of sources and dilutions of antibodies used for histo/immunochemistry in this study.

| Antibody table |  |  |  |
| --- | --- | --- | --- |
| Antibody | Species | Dilution | Company (Catalog number) |
| GFP | Chicken | 1:500 | Aves (GFP-1020) |
| Sox9 | Goat | 1:500 | R&D Systems (AF3075) |
| GFAP | Rabbit | 1:500 | Millipore (AB5804) |
| CNPase | Mouse | 1:200 | Millipore (MAB326R) |
| Tuj1 | Mouse | 1:500 | Covance (MMS435P) |
| Neurocan | Mouse | 1:500 | Millipore (MAB5212) |
| DCX | Goat | 1:500 | Santa Cruz Biotechnology (sc-8066) |
| NeuN | Mouse | 1:500 | Millipore (MAB377) |
| PDGFrβ | Rabbit | 1:500 | Abcam (ab32570) |
| Click-it$\text{\textcircled{C}}$ Plus EdU | | | Molecular probes by Life (C10638) |
| DAPI |  | 1:1000 | Sigma (D9542) |
| Anti-rabbit cy3 secondary antibody | Donkey | 1:500 | Jackson Immuno Reasearch (711-166-152) |
| Anti-mouse cy3 secondary antibody | Donkey | 1:500 | Jackson Immuno Reasearch (715-165-140) |
| Anti-chicken Alexa 488 secondary antibody | Donkey | 1:500 | Jackson Immuno Reasearch (703-545-155) |

### Cell culture analysis

The phenotypic characterization was observed and photographed (×200) with MetaMorph Software (Leica).

For the neurosphere proliferation assay, neurospheres were sampled from one culture dish in quadruplicate (250 ml per time) and the numbers of neurospheres were counted (Figure 1K-N). The total number of neurospheres was calculated per total volume of media (30–35 ml per dish). For the neurosphere differentiation assay and primary bOEC or fibroblast cultures characterization (Figures 2O-S, 1B and C and 4B and C respectively), we quantified the marker expression by counting at least 6 fields of views selected in a randomized fashion (20× magnification) per well under fluorescence microscope. As previously described, primary bOEC cultures is composed of around 20-30% of bOECs and 70% of fibroblasts (Figure 1B and C).

At least two wells for one marker per animal and condition were used for quantification of a minimum of 3 animals per group. The total number of cells was obtained counting DAPI+ nuclei stained.

### Immunohistochemistry

Animals were deeply anesthetized with sodium pentobarbital (150 mg/kg body weight) and perfused transcardially with PBS followed by ice-cold 4% formaldehyde in PBS. Dissected spinal cords were post-fixed overnight at 4°C and cryoprotected in 30% sucrose solution in PBS. The tissues were embedded into OCT (Tissue-Tek^®^, Sakura). Coronal and sagittal 20µm sections were collected alternating on ten (8-10 sections per slide). Sections were incubated with blocking solution (10 % Normal Donkey serum, 0.3% Triton-X100), then incubated overnight at room temperature in a humidified chamber with primary antibodies diluted in blocking solution. The following antibodies were used (Table 1). EdU was detected with the Click-iT® EdU Alexa Fluor® 594 imaging kit (Invitrogen) using the Manufacturer’s instructions. After washing, antibody staining was revealed using species-specific fluorophore-conjugated (Cy3, Cy5 or Alexa 488 from Jackson Immuno Research) secondary antibodies. Sections were counterstained with DAPI (1µg/mL) and sections were coverslipped with Vectashield mounting media (Vector Labs). Pictures were taken using a Zeiss LMS700 microscope. Image processing and assembly was performed in ImageJ or Photoshop.

### Image acquisition analysis

Confocal representative images of the lesion site and spinal cords were acquired using the Zeiss LSM700 or Zeiss LSM800 microscope set up.

SCI and transplantation studies used coronal sections for analysis. For cell culture analysis, 6 - 8 randomly selected views per well were used for analysis, and at least 3 wells per animal were used for statistical analysis. Quantification of the number of cells were performed using the Zeiss Apotome2 microscope set up. The quantification of cells was performed in 2-4 sections per animal. For each experimental group and staining, 3-9 animals were analyzed.

### Real time PCR

Real time PCR experiments were performed to evaluate the level of mRNA expression of the axonal growth inhibitory and axonal growth permissive molecules and also neurotrophic factors. Mice were killed 2 weeks after SCI and spinal cord were immediately dissected on dry ice. Total RNAs from these tissues samples were extracted using Tri-reagent (Sigma) and Nucleospin RNAII kit (Macherey-Nagel) according to the manufacturer’s protocol. From each sample, 1.5 μg of total RNA was converted into single stranded cDNA using the ImPromII reverse transcriptase kit (Promega) with random primers (0.5μg/ml). Real time PCR experiments were performed and monitored by ABI Prism 7500 Sequence Detection System (Life Technologies). The primer pairs used for the different gene analyses have been previously described (Anderson et al., 2016; Sabelstrom et al., 2013). Mouse glyceraldehydes-3-phosphate dehydrogenase (GAPDH) cDNA was used as control. Relative expression between a given sample and a reference sample was calculated using the 2^-ΔCt^ method, where ΔCt is the difference in the Ct values for the target gene and the reference gene. All experiments were performed with two technical replicas.

### Statistical analysis

All data are presented as mean ± standard deviation (SD). Statistical analyses were conducted using GraphPad Prism software, version for windows (GraphPad Software; 8.0.1.244). Statistic were run with Student’s T-test for comparing two groups and Student’s T-test with Bonferroni’s correction for more than two groups’ comparisons. Detailed statistical analyses for each assay are provided in the figure legends. \**P*<0.05, \*\**P*<0.01, \*\*\**P*<0.001 and #*P*<0.0001. For each experimental group and staining, 3-9 animals minimum were analyzed.

## RESULTS

### Transplantation of bOECs increases ependymal cell proliferation *in vivo* and *in vitro*

Before studied the effects of transplanted bOEC on tissue repair after SCI, we first evaluated the survival of bOECs in WT C57BL/6 mice. To do so, we isolated bOECs from luciferase animals (Cao et al., 2004; Sheikh et al., 2007) and transplanted the luciferase+ bOECs into WT animals after SCI. As previously described by other teams, the number of transplanted bOECs decreased quickly and significantly after 2 weeks post injury in comparison to one-week post SCI (Figure 1D and E).

To assess how bOECs could promote recovery after SCI although their survival timeframe is short after transplantation, we hypothesized that bOECs could affect endogenous cell types, especially stem and progenitor cells that can contribute to tissue repair after SCI. Ependymal cells around central canal rarely divide in intact spinal cord but start to proliferate vigorously after SCI, and can be lineage traced by FoxJ1-CreER mice (Barnabé-Heider et al., 2010; Meletis et al., 2008; Rodrigo Albors et al., 2023)). First, after permanently labeling ependymal cells, we confirmed that bOEC transplantation did not alter YFP recombination in these cells, either in the intact or injured spinal cord (Figure 1F). These findings are consistent with previous studies (Figure 1F) (Barnabé-Heider et al., 2010; Li et al., 2016; Meletis et al., 2008). Next, to investigate whether bOEC transplantation affects ependymal cell proliferation both *in vivo* and *in vitro* after SCI, we transplanted bOECs into FoxJ1-CreER^T2^-YFP mice and labeled the proliferative cells with EdU administration via drinking water during 1 week. We found that 2 weeks after SCI, there was a significant increase of EdU incorporation in ependymal lineage cells in the bOEC group compared to vehicle (injection of media, “DF10S”) (83,07 % ± 0,913% versus 72,5 ± 1,745%) (Figure 1G and H), suggesting that bOEC transplantations after SCI enhance the proliferative potential of neural stem cells (i.e. ependymal cells) *in vivo*.

We then assessed the spinal cord self-renewal potential *in vitro* after OEC transplantation. To do so, we used the same surgical protocol as described above and performed neurosphere assay in FoxJ1-CreER^T2^-YFP. Seven days after SCI, dissociated spinal cords were cultured to produce neurospheres (Figure 1I). We found that the stem cell potential of spinal cord was still only confined in ependymal cells with bOEC transplantation, suggesting that OECs did not promote another stem cell niche with or without SCI. This is supported by the *in vitro* recombination pattern in the primary neurospheres, which was identical between OEC groups with or without SCI (Figure 1J). Moreover, during passages, 100% of neurospheres were YFP positive, confirming that all stem cell potential was only confined to ependymal cells upon bOEC transplantation (Figure 1J). These recombination patterns in primary and further generations of neurospheres were similar to the ones previously described in uninjured and injured animals without bOEC transplantation (Barnabe-Heider et al., 2010; Li et al., 2016).

To further study the effects of bOECs on self-renewal potential of ependymal cells *in vitro*, we performed neurosphere assay under four conditions, including 1) no injury as baseline, 2) SCI with DF10S, 3) no injury but with bOEC transplantation and 4) SCI with bOEC transplantation. We quantified and compared the total number of generated neurospheres per 100,000 cells for each passage and set the group with no injury nor bOECs as one-fold (Figure 1K-N). We found that SCI resulted in two times more neurospheres than healthy control, while bOEC transplantation after SCI can lead to double amount of neurospheres compared to SCI group. Interestingly, without SCI, the self-renewal potential of ependymal cells was also able to be activated highly as the SCI group, suggesting that either SCI or bOEC transplantation is sufficient to increase the self-renewal potential, while the effect reaches the highest level after both SCI and bOEC transplantation (Figure 1K). These primary neurospheres were dissociated to generate secondary neurospheres. In this context, the non-SCI and SCI + DF10S groups appeared similar, whereas the transplanted groups (non-SCI + bOECs and SCI + bOECs) generated twice as many neurospheres (Figure 1L). Subsequently, as previously described, self-renewal potential gradually becomes constant over passages regardless of the paradigm tested (Figure 1M and N) (Li et al., 2016).

Therefore, these *in vivo* and *in vitro* data indicate that bOEC transplantations in spinal cord can highly enhance the self-renewal potential of ependymal cells.

### bOEC transplantation changes ependymal cell progeny

It has been previously shown that after SCI, proliferative ependymal cells leave the central canal, migrate to the lesion site and differentiate into glial cell lineage (Barnabe-Heider et al., 2010; Meletis et al., 2008; Sabelstrom et al., 2013). We further investigated the effects of bOEC transplantation on the fate of ependymal progeny after spinal cord dorsal hemisection (T9-T10). FoxJ1-CreER^T2^-YFP mice were used to fate map the ependymal cells and follow the activation, migration and differentiation of their progeny after SCI *in vivo* (Figure 2A-N). Two weeks and fourteen weeks after surgery, we sacrificed the animals and performed immunohistochemistry analysis. We found that 2 weeks after SCI, ependymal cells start to leave the central canal and migrate to the lesion site in both DF10S and bOEC transplantation groups (Figure 2A and E). Similarly to previous studies (Barnabe-Heider et al., 2010), we observed that the ependymal progeny highly express Sox9 in both groups at around 80%, with a slight but significant increase in SCI + bOECs group (Figure 2C, G and J). Our analyses show also that bOEC transplantations did not change the number of YFP+/Olig2+ cells 2 weeks after SCI compared to DF10S group (Figure 2D, H and K). However, while it was consistent with previous study that ependymal lineage gives rise to around 15%-20% GFAP+ cells (Barnabe-Heider et al., 2010), we found that there was almost three times more YFP+/GFAP+ cells in bOECs group than DF10S group (45.0 ± 4.5% V.S. 15.5 ± 1.9%), suggesting that bOECs induce more ependymal cells to differentiate into astrocytes, but not more into oligodendrocytes (Figure 2B, F and I). After 14 weeks, a higher proportion of YFP+/Sox9+ cells was still observed in transplanted group in comparison to DF10S group while no significance was observed in YFP+/GFAP+ cells (Figure 2L-N). These data suggest that bOEC transplantation strongly induces an astrocytic differentiation of ependymal cells *in vivo* at early stage after SCI.

To further investigate whether bOEC transplantation affects stem cell differentiation *in vitro*, we performed differentiation assay on the neurospheres derived from ependymal cells as previously described (Meletis et al., 2008 and Figure 1I-N). Neurospheres from spinal cord display multipotency and can give rise to astrocytes, oligodendrocytes and neurons *in vitro* (Barnabe-Heider et al., 2010; Li et al., 2018a; Li et al., 2016). After differentiation of the primary neurospheres, we used antibody CNPase to identify oligodendrocyte lineage and Tuj1 for neuronal lineage respectively (Figure 2O-S). Similarly to previous studies (Li et al., 2018b; Li et al., 2016), adult spinal cord neurospheres generate a low proportion of CNPase+ cells at around 0.5% (Figure 2O and R), but this capacity was significantly increased by two fold after SCI (Figure 2R), regardless of bOEC transplantations. On the contrary, bOEC transplantation significantly increased the proportion of Tuj1+ cells compare to the non-transplanted groups in both uninjured conditions (from 1.3 ± 0.06% to 3.6 ± 0.61%) and injured conditions (from 2.3 ± 0.17% to 3.8 ± 0.23%), respectively (Figure 2Q and S) while no significance was observed for the proportion of Sox9+ cells (Figure 2P and Data not shown). These data suggest that bOEC transplantation increases neuronal differentiation while SCI increases oligodendrocytic differentiation of ependymal cells *in vitro*.

### bOEC transplantation enriches the scar environment for axonal regrowth and promotes neuronal survival after SCI

It has been suggested that after SCI, astrocytes become reactive and contribute to the inhibitory glia scar by assembling and secreting a large amount of CSPG (Chondroitin Sulfate Proteoglycan) that act as a major barrier for regenerating axons (Cregg et al., 2014; O’Shea et al., 2017). However, astrocytes derived from ependymal cells express low level of this axonal growth-inhibiting CSPG (Meletis et al., 2008). As bOEC transplantation increases the differentiation of ependymal cells into astrocytes (Figure 2I), we further characterized how such enhanced astrocytic differentiation could affect the scar environment. First, we analyzed the expression of Neurocan, one major inhibitory produced CSPG upon SCI (Jones et al., 2003). Two weeks after SCI, we found that some YFP+/GFAP+ cells in SCI + DF10S control group expressed Neurocan, while bOEC transplantation group had almost none expression in the lesion core (Figure 3A and B), suggesting that the induced astrocyte differentiation by bOECs had low expression of Neurocan. To further study more broadly whether bOEC transplantation could modulate the scar formation by inducing a microenvironment with higher axonal regrowth potential, we performed qPCR experiments for a number of axonal growth permissive and axonal growth inhibitory molecules, using the similar list reported by Anderson et al., 2016 (Figure 3C and D, Table 2). We took the lesion tissues 2 weeks after SCI with and without bOEC transplantation to perform qRT-PCR, and compared these two groups to uninjured WT animals used as the baseline control (indicated as dashed line in Figure 3C and D) (Table 2). Similarly to previous study, we found in both groups, independently of bOEC transplantation, that SCI induced higher gene expression related to axonal growth (Anderson et al., 2016). Even though bOEC transplantation induced a decrease or an increase in the expression of a small number of the axonal growth permissive genes, 5 out of 10 from the top 10 axonal growth permissive genes in this category were found unchanged with or without bOEC transplantation (Figure 3C). However, 9 out of 10 genes from the top 10 axonal growth inhibitory genes expressed by the two groups were significantly down regulated in bOEC group (Figure 3D). These results suggest that bOEC transplantation enriches the microenvironment of the injured spinal cord for better axonal regrowth, partly by inhibiting the expression of axonal growth inhibitory molecules.

**Table 2.**
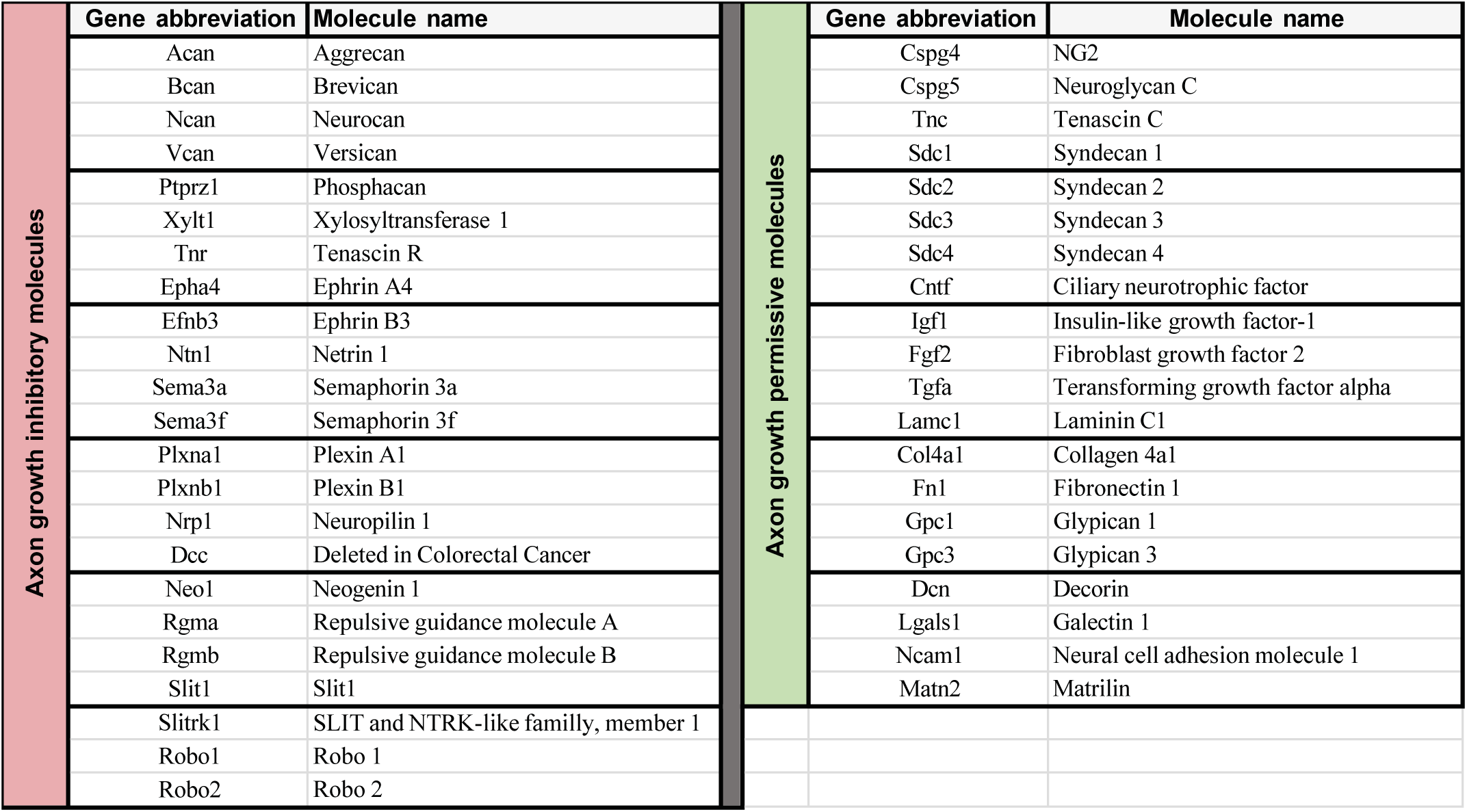
Table of genes for axonal growth inhibitory and axonal growth permissive molecules analyzed by qPCR. This table lists the genes abbreviation and full name of 43 axon-growth-modulating molecules for which gene expression levels were analyzed in this study (Anderson et al., 2016; Sabelstrom et al., 2013).

As ependymal cells can be modulated by bOECs for better regenerative potential, and previous study showed that ependymal cells secret neurotrophic factors to maintain neuronal survival after SCI (Sabelstrom et al., 2013), we further analyzed whether the increased number of ependymal cells by bOECs could promote better neuronal survival. Our data showed that the number of NeuN+ cells decreased over time (from 2 to14 weeks after SCI) in the SCI + DF10S group, while the number of surviving neurons was increased at 2 and 14 weeks after SCI in bOEC transplanted group (Figure 3E-H), suggesting that bOEC transplantation increased neuronal survival.

### The effects of cell transplantation on ependymal cells are bOEC-dependent

As previously described, primary bOEC cultures consist of a mixed population of fibroblasts and bOECs (Figure 1B and C). We therefore aimed to address whether the effects we observed are solely contributed by bOECs, or could be from the fibroblasts in the culture. To this end, we obtained purified olfactory fibroblast cultures and transplanted them into FoxJ1 mice following SCI (Figure 4A). This approach enabled us to obtain purified fibroblast cultures, with a PDGFRβ-positive cell purity of nearly 100% (Figure 4B and C). These cultures were then transplanted into FoxJ1-CreER^T2^-TdTomato mice in order to assess ependymal cell proliferation and differentiation two weeks after SCI. As a first step, to ensure that the recombination rate of EdU⁺ cells in FoxJ1⁺ ependymal cells was not affected by performing the experiments in a TdTomato reporter line, we measured the proliferation of ependymal cells around the central canal (Figure 4D). We demonstrated that the recombination rate of TdTomato⁺/EdU⁺ (TdTom⁺) cells was comparable to that observed in Figure 1H, and was increased in mice transplanted with bOECs compared to DF10S-treated mice (Figure 4D). Then, we investigated the effects of fibroblast transplantation (Figure 4D-L). Unlike bOEC transplantation, fibroblast transplantation did not alter the proportion of TdTomato⁺/EdU⁺ cells, which remained comparable to that observed in the SCI + DF10S control group (Figure 4D and E). Similarly, fibroblast transplantation had no effect on the differentiation of ependymal cells (Figure 4F–K), and did not change the proportion of TdTomato⁺/GFAP⁺, TdTomato⁺/Sox9⁺, or TdTomato⁺/Olig2⁺ cells, in contrast to bOEC transplantation (Figure 4F, H, and J, respectively). Finally, we assessed the effect of fibroblast transplantation on neuronal survival after SCI and found that it did not modulate neuronal survival compared to the SCI + DF10S control group (Figure 4L).

Altogether, these results indicate that the effects of cell transplantation on ependymal cells after SCI are specifically dependent on bOECs.

### The endogenous ependymal cells are required for regenerative effects promoted by bOEC transplantation

While we have shown that ependymal cells are positively modulated by the transplantation of primary bOECs, it is still unclear if transplanted bOECs affect different cell types directly or through ependymal cells as an intermediate modulator. As ependymal cells were found to be vital for neuronal survival by modulating the microenvironment after SCI (Sabelstrom et al., 2013), we investigated whether the lack of proliferative ependymal cells would change the effects induced by transplanted bOECs. To this end, we took advantage of another transgenic mouse model FoxJ1-Rasless to specifically ablate the proliferation of ependymal cells and thereby block the generation of ependymal cell progeny upon injury (Sabelstrom et al., 2013). First, we transplanted the same number of bOECs into FoxJ1-Rasless animals as described above (Figure 5A) and investigated the effects of ependymal cell proliferation on bOEC survival after SCI using bioluminescence as described previously (Figure 1D and E). As described in Figure 1, the number of transplanted bOECs decreases quickly at 1- and 2-weeks post SCI (Figure 5B).

To study the role of bOEC transplantation on spinal cord microenvironment, we used the same list of axonal growth permissive and axonal growth inhibitory genes as shown in Figure 3C and D. Using FoxJ1-Rasless animals 2 weeks after SCI with and without bOEC transplantations for qPCR, we found that bOEC transplantation failed to modulate axonal growth inhibitory genes (Figure 5C and D), differently from that in FoxJ1-CreER^T2^-YFP mice (Figure 3D), suggesting that ependymal cell proliferation and reactivity are essential for establishing a non-inhibitory environment for axonal regrowth in the context of bOEC transplantation.

Moreover, we investigated whether the neuronal protection on mature neurons NeuN+ effects by bOEC transplantation is also dependent on ependymal cells. We used both FoxJ1-CreER^T2^-YFP and FoxJ1-Rasless animals for SCI and bOEC transplantations at 2 and 14 weeks after SCI, and we compared the neuronal survival levels. We found that 2 weeks after SCI, bOEC transplantation achieved neuronal protection more significantly than DF10S group after SCI in FoxJ1-CreER^T2^-YFP mice, but failed to do so in the absence of proliferative ependymal cells in FoxJ1-Rasless mice where the cell cycle of ependymal cells is blocked (Figure 5E). We also found the similar consequence on neuronal survival 14 weeks after SCI and transplantation (Figure 5F). These data suggest that ependymal cells are required for neuronal protection, and play an essential role in promoting neuronal survival and microenvironment modulation by bOEC transplantation.

The adult spinal cord has long been considered as a non-neurogenic tissue. Indeed, no generation of mature neurons occurs in this tissue under normal conditions or after injury. However, it has been shown that immature DCX⁺ neurons can be generated in the adult spinal cord (Habib et al., 2016). Therefore, we assessed the density of immature DCX⁺ neurons in FoxJ1-CreER^T2^-YFP and FoxJ1-Rasless mice two weeks after SCI and bOEC transplantation. These analyses revealed that the density of DCX⁺ cells was markedly reduced in FoxJ1-Rasless mice and is identical to SCI + DF10S ones (Figure 5G, see dash line for SCI + DF10S group).

## DISCUSSION

It has been shown in both animal and human studies that OECs can promote regeneration after SCI, but the mechanism is largely understudied. In this study, we demonstrate that activated ependymal cells are essential mediators of the regenerative effects induced by bOEC transplantation. Using various transgenic mouse models, we found that: i) bOEC transplantation enhances the stem cell potential of spinal cord progenitors following injury (Figure 1); ii) bOECs modulate the differentiation capacity of ependymal cells, thereby reshaping the lesioned microenvironment and promoting neuronal survival (Figures 2 and 3); iii) the observed effects are bOEC-dependent and cannot be replicated by fibroblast transplantation (Figure 4) and iiii) the beneficial effects of bOEC transplantation on the injured spinal cord are dependent on the activation of endogenous spinal cord stem cells (Figure 5).

OEC transplantation has been widely recognized as a promising therapeutic strategy for SCI, as demonstrated in both animal models and clinical trials (Delarue and Guérout, 2022; Phelps et al., 2025). These studies have shown that OECs secrete factors that promote axonal regrowth and improve functional recovery (Kubasak et al., 2008; Lakatos et al., 2000; Li et al., 1997; Polentes et al., 2004; Ramón-Cueto et al., 2000; Stamegna et al., 2011; Tabakow et al., 2013). However, OECs typically survive only 2 to 4 weeks after transplantation, whereas functional recovery can persist for months (Khankan et al., 2016; Reshamwala et al., 2019). This raises the question of whether the observed long-term benefits result directly from the transplanted OECs or from their capacity to modulate endogenous cell populations that sustain regenerative processes.

Previous findings have identified ependymal cells surrounding the central canal as the primary stem cell population of the adult spinal cord. These cells have been shown to be essential for glial scar formation and for supporting neuronal survival after SCI (Barnabé-Heider et al., 2010; Rodrigo Albors et al., 2023; Sabelström et al., 2013).

In our study, we used FoxJ1-CreER^T2^ mice to lineage-trace ependymal cells and investigate their behavior following bOEC transplantation. We found that bOECs enhance both the proliferation and astrocytic differentiation of ependymal cells *in vivo*, while also promoting their self-renewal capacity *in vitro* (Figures 1 and 2). These findings suggest that bOECs modulate ependymal cell activity, contributing to a more structured and possibly beneficial glial scar. Importantly, these results provide a potential explanation for the long-lasting regenerative effects observed after OEC transplantation, even after the transplanted cells themselves have disappeared.

A key concern, however, is whether the increased number of astrocytes derived from ependymal cells might inhibit axonal regrowth, since the glial scar—largely composed of astrocytes—has historically been viewed as a major barrier to regeneration. (Anderson et al., 2014; Burda and Sofroniew, 2014). While it is true that astrocytes in the scar can express molecules that inhibit axonal growth, several studies have demonstrated that they also release growth-promoting factors. Moreover, depleting astrocytes, including those derived from ependymal cells, has been shown to exacerbate injury and impair recovery (Anderson et al., 2016; Sabelström et al., 2013). In our study, we observed an increase in ependymal cell-derived astrocytes following bOEC transplantation. Notably, these astrocytes did not express neurocan, a well-known inhibitory CSPG, suggesting that this specific population may not suppress—but rather support—axonal regeneration (Figure 3). As such, the enhanced generation of astrocytes from ependymal cells induced by bOECs could contribute positively to spinal cord repair and structural reinforcement after injury.

Based on the previously reported list of genes associated with permissive and inhibitory axonal regrowth molecules (Anderson et al., 2016; Delarue et al., 2019), our qPCR analysis revealed that the expression of most inhibitory axonal growth genes was significantly reduced following bOEC transplantation (Figure 3). However, when using Rasless mice, in which ependymal cell proliferation is specifically blocked, we did not observe notable differences in the expression profiles of these genes between the DF10S control and the bOEC-transplanted groups after SCI (Figure 5). This suggests that the presence of proliferating ependymal cells is necessary for bOECs to effectively modulate the microenvironment in favor of axonal regeneration.

Moreover, previous studies have shown that ependymal cells are essential for neuronal survival following SCI, particularly through their regulation of neurotrophic factor expression (Sabelström et al., 2013). In our experiments, we observed a greater number of surviving neurons in the bOEC-transplanted group at both 2- and 14-weeks post-injury, likely due to the expansion of the ependymal cell population induced by the transplantation (Figure 3). However, this neuroprotective effect was only evident in the presence of proliferative ependymal cells. In Rasless mice lacking ependymal cell proliferation, no significant differences in neuronal survival were found between the bOEC and DF10S groups, indicating that the beneficial effects of bOEC transplantation on neuronal survival are dependent on the presence and activation of ependymal cells (Figure 5).

Since primary cultures of bOECs consist of a mixed population of fibroblasts and bOECs, it is plausible that the effects of transplantation on ependymal cells are driven by the transplantation procedure itself rather than by the intrinsic properties of the OECs. To determine whether these effects are specifically dependent on bOECs, we generated purified cultures of fibroblasts, which were then transplanted into SCI mice and compared to mice that received bOEC transplants. This approach allowed us to demonstrate that the previously observed effects on ependymal cells are indeed OEC-dependent. Specifically, fibroblast transplantation did not affect the proliferation or differentiation of ependymal cells *in vivo* (Figure 4).

Proof-of-concept studies have shown that modulating the reactivity of specific cellular populations through genetic manipulation or viral delivery can promote both tissue repair and functional recovery (Anderson et al., 2018; Dias et al., 2018; Llorens-Bobadilla et al., 2020; Squair et al., 2023). However, such approaches are not currently applicable in a clinical setting. In contrast, cell transplantation offers a more clinically feasible strategy by modulating the spinal cord environment through exogenous intervention. Nevertheless, its effects appear to be limited, likely due to the short survival time of the majority of transplanted cells, which typically persist for only a few days to weeks (Reshamwala et al., 2019). Our study highlights that these two seemingly opposing strategies—genetic modulation and cell transplantation— are not mutually exclusive but, in fact, complementary. We demonstrate that the transplantation of differentiated cells, which do not integrate into the spinal cord parenchyma and exhibit limited post-transplant survival, can still exert long-lasting modulatory effects on the injured spinal cord without posing a risk of tumor formation.

Taken together, our study reveals for the first time that bOEC transplantation activates endogenous ependymal cells, which in turn contribute to the enhancement of the injury microenvironment and support neuronal survival after SCI. Our findings propose a model in which ependymal cells play a central role in mediating the beneficial effects of cell transplantation-based therapies for SCI. This work opens new avenues in the field of regenerative medicine, emphasizing the therapeutic potential of activating, recruiting, and modulating endogenous spinal cord stem cells as a strategy for future SCI treatment.

## ACKNOWLEDGEMENTS

We are grateful to Dr. Jonas Frisén for sharing the FoxJ1-CreER^T2^ and FoxJ1-Rasless transgenic mouse lines as well as critical comments.

## AUTHOR CONTRIBUTIONS

N.G. conceptualized the project.

Q.D., A.H., X.L. and N.G. designed the experiments.

Q.D., A.H., C.C., M.D.G. and X.L. performed the experiments.

A.H., X.L. and N.G. analyzed the results.

D.V. provided reagents, techniques and scientific input.

Q.D., X.L. and N.G. wrote the article.

